# Climatic seasonality and topographic complexity shape plant growth form distributions in the Canary Islands

**DOI:** 10.1101/2025.09.19.677327

**Authors:** Lucas S. Jansen, Ryan F.A. Brewer, Juli Caujapé-Castells, Luis Valente, Frederic Lens

## Abstract

Plant growth forms reflect the environmental conditions under which they evolved and persist. Yet, previous studies linking environmental conditions with plant growth form have overlooked that different growth form types arose through distinct evolutionary pathways. For instance, some woody angiosperm lineages evolved woodiness from a herbaceous ancestor, whereas others have retained their woodiness throughout evolutionary history. The Canary Islands are a global hotspot of insular woodiness (woodiness trait evolved on the islands from herbaceous colonisers), raising the question of whether insular woody species generally occur in different environmental conditions than other growth forms on the islands. Here, we use a novel pipeline to extract, filter, and analyse publicly available spatial data to identify the environmental correlates of plant growth forms in the angiosperm flora of the Canary Islands, accounting for their distinct evolutionary histories as well as differences in life span. To assess whether and how five growth forms (annual herbaceous, perennial herbaceous, ancestral woodiness, derived woodiness, and insular woodiness) differ in environmental niche space, we applied a phylogenetic principal component analysis to a dataset of ∼1,000 native Canary Island angiosperm species. We show the herbaceous and woody growth forms have an unexpectedly high overlap in niche space. The subtle differences are explained mostly by climatic seasonality (in both temperature and precipitation) and topographic complexity. We use phylogenetic ANOVAs to identify significant pairwise differences between the five growth forms, and find significant differentiation between five environmental variables, with differences between a combined woody and a combined herbaceous niche, and between the different evolutionary histories across the woody growth forms. Through our approach, the collection and analysis of publicly available spatial and environmental data can be automated, facilitating research into the environmental niche.

## 1 Introduction

Since transitioning from water to land ∼500 million years ago (Morris et al., 2018), terrestrial plant life has evolved into a diverse array of growth forms: from the structurally simple extinct early land plants to the complex forms of extant herbaceous and woody seed plants (Stewart and Rothwell, 1993). This myriad of growth forms reflects the wide range of environmental conditions and biological interactions that have given rise to the extant global flora (Rowe & Speck, 2005). Yet, despite centuries of study (von Humboldt, 1807), much remains unknown about the relationship between environmental factors and different forms of plant life.

Many different ways to classify growth forms exist, from the classification system of Raunkiaer (e.g., chamaephyte, phanerophyte; Raunkiaer, 1934; Irl et al., 2020), to a simple binary trait state describing the aboveground stem (herbaceous vs woody; Poppenwimer et al., 2023), a more complex multistate that takes into account life span (annuality vs perenniality) and evolutionary history (Hooft van Huysduynen et al., 2021), or based on descriptions of the entire plant (e.g., trees, shrubs, epiphytes, terrestrial herbs; Taylor et al., 2023). Here, we adopt the multistate classification to look into the relationship with environmental factors. This relationship is clearly visible. For example, at the global scale, a recent study found that the extant distributions of annual herbaceous species — i.e., non-woody species that complete their life cycle in one year — are driven by prolonged high temperatures and seasonal drought (Poppenwimer et al., 2023), with these species surviving the harshest periods of the year as seeds. In contrast, perennial species (those with longer-lived belowground and/or aboveground organs that are either herbaceous or woody) are more frequently encountered in tropical conditions and at the highest latitudes, creating a non-linear global latitudinal relationship in the distribution of annuals and perennials (Friedman, 2020; Poppenwimer et al., 2023; Taylor et al., 2023).

However, an often overlooked factor in macroecological studies of plant growth form is that extant growth forms have evolved through different evolutionary pathways; by grouping growth forms with different evolutionary origins, based on extant phenotype, their true environmental correlates may be obscured. For example, because angiosperms originated from woody gymnosperm-like ancestors, it is generally accepted that the most recent common ancestor of angiosperms was woody, giving rise to today’s “ancestrally woody” species that retained their growth form throughout evolutionary history (Gerriene et al., 2011; Doyle, 2012). Many woody angiosperm lineages later independently lost woodiness, evolving into (annual or perennial) herbaceous plant taxa, accounting for nearly 50% of extant angiosperm species (Klimeš et al., 2022). Notably, multiple herbaceous lineages have re-evolved woodiness, an evolutionary reversal known as phylogenetically derived woodiness (Zizka et al., 2022). However, despite the massive number of transitions from herbaceousness towards derived woodiness in angiosperms (Zizka et al., 2022), the majority of angiosperm species we see today are ancestrally woody, i.e. have retained woodiness from woody ancestors (Hooft van Huysduynen et al., 2021).

Within derived woody growth types, a further distinction based on geographical context can be made. “Insular woodiness” refers to when phylogenetic inference shows that woodiness evolution occurred on an island after being colonised by a herbaceous continental species (Carlquist, 1974; Zizka et al., 2022). Insular woodiness differs from (continental-)derived woodiness, which are island species which shifted from herbaceousness to woodiness on a continent before colonising the island. This distinction recognises insular woodiness as a typical island phenomenon, considered part of the classic island syndrome set of traits (Darwin 1859; Carlquist, 1974; Lens et al., 2013a; Zizka et al., 2022). Interestingly, many iconic island lineages are insular woody, such as the Hawaiian silverswords (*Argyroxiphium, Dubautia, Wilkesia*, Asteraceae; Baldwin & Sanderson, 1998) and Canary Island viper buglosses (*Echium*, Boraginaceae; Böhle et al., 1996).

Several hypotheses have been proposed to explain the evolution of insular woodiness, such as competition for capturing sunlight in dense herbaceous populations growing in open environments (Darwin, 1859), longer life spans due to a lack of large native herbivores on islands (Carlquist, 1974), or due to a favourable aseasonal insular climate (Carlquist, 1974). More recently, a theory proposed by Lens et al. (2013b), stated that drought could have been an important global driver of woodiness evolution. Experimental evidence for this theory, provided by, among others, Dória et al. (2018), found that stems of insular woody daisies (*Argyranthemum*, Asteraceae) were better able to avoid drought-induced gas bubbles in their water-conducting vessels compared to their herbaceous relatives. This drought hypothesis was further supported by a global macroecological analysis (Zizka et al. 2022), which found that native insular woody species occurrence is correlated with drought-related variables, although variables related to favourable climate (only for oceanic islands) and lack of herbivory (for oceanic islands and continental fragments) were also correlated.

Research on the macroecology of growth form in island plants has hitherto overlooked the distinct evolutionary histories associated with these growth forms (such as insular woodiness, ancestral woodiness, and derived woodiness). Consequently, if we aim to disentangle the relationship between environment and growth form on islands, we need comprehensive analyses addressing fine-scale environmental correlates and considering the different evolutionary origins of growth forms (Sauquet & Magallón, 2018). Here we perform such an analysis, focusing on the Canary Islands. The Canary Islands archipelago has one of the most diverse and best-studied insular floras globally and is a hotspot for insular woodiness, in both numbers of species and shifts towards insular woodiness (Beierkuhnlein et al., 2021; Zizka et al., 2022; Barajas Barbosa et al., 2023). To investigate fine-scale processes underlying plant growth forms, we categorised the non-monocot angiosperm flora of the Canary Islands into two herbaceous (annual and perennial herbaceous) and three woody (ancestral, derived, and insular woodiness) growth forms. We do not include monocots as they never develop woodiness due to the lack of a vascular cambium (Carlquist, 2012; Zizka et al., 2022).

We test the following hypotheses: (1) due to different eco-physiological requirements, woody and herbaceous species on the Canaries occur in localities with different environmental characteristics, detectable through differences in niche space. Within the two herbaceous and three woody growth forms, we expect additional differences in niche space due to differences in life span and evolutionary history, respectively. (2) As insular woody lineages are, by definition, the only ones that evolved woodiness *in situ*, we expect the insular woody niche to deviate most from that occupied by herbaceous growth forms, reflecting a possible adaptive benefit of this shift in growth form on the islands. (3) In line with previous studies addressing the drivers of insular woodiness (Lens et al., 2013a; Hooft van Huysduynen et al., 2021), we expect that variables related to aridity and aseasonality impact the niche of insular woody species (Carlquist, 1974; Zizka et al., 2022).

To assess these hypotheses, we follow four steps. First, we develop a novel bioinformatics pipeline to extract environmental variables based on open-source plant occurrence data and use it to compile a dataset of environmental variables for the non-monocot angiosperm Canarian flora. Secondly, we use a phylogenetic principal component analysis (Revell, 2009) to disentangle the variables that delineate environmental niche space for the angiosperm flora of the Canaries, identifying the environmental correlates for each growth form whilst accounting for phylogenetic ancestry. Third, we apply phylogenetic ANOVAs to test for pairwise differences in the association of each variable with the distribution of each growth form. Finally, we highlight the impact of a variable that was most significant in the phylogenetic ANOVAs: rugosity, i.e. topographic complexity, measured using a terrain ruggedness index.

## 2 Methods

### 2.1 Locality data extraction

#### 2.1.1 Study region & Target species

The Canary Islands are a volcanic archipelago ∼100km to the west of southern Morocco and consist of seven main islands (El Hierro, La Palma, La Gomera, Tenerife, Gran Canaria, Fuerteventura and Lanzarote) and two small islands (<30 km2) off the coast of Lanzarote (La Graciosa and Alegranza; Figure 1). The Canarian archipelago has one of the most species-rich oceanic island floras globally, as well as the highest number of insular woody species globally (Triantis et al. 2015; Zizka et al., 2022).

**Figure 1.**
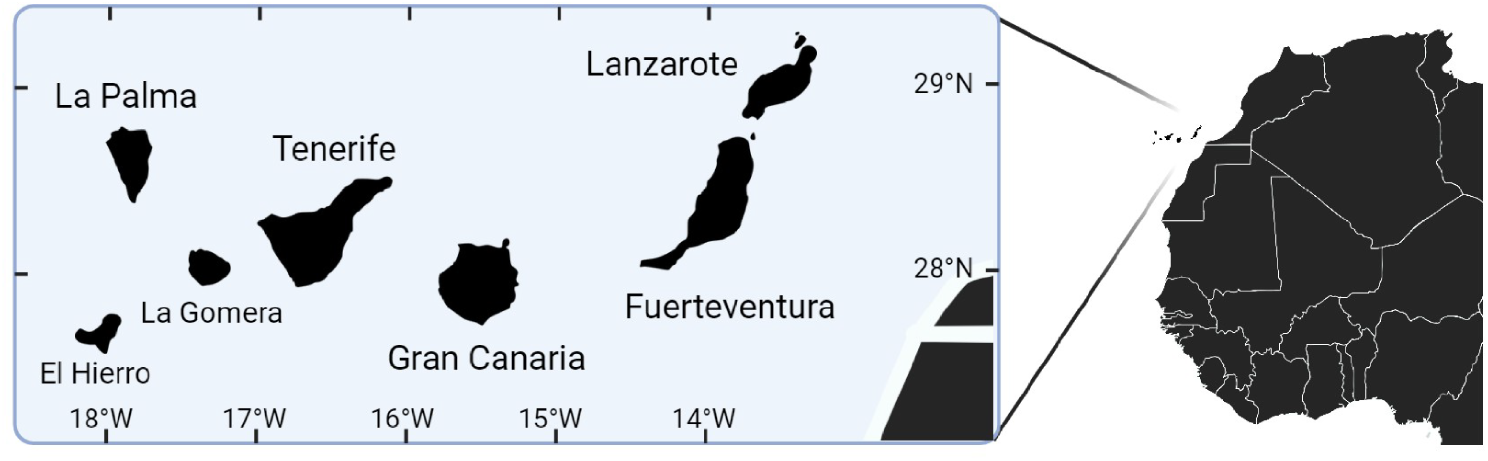
Map of the Canary Islands archipelago. The names of the seven main islands are included, as well as latitude and longitude (degrees) ticks.

**Figure 2.**
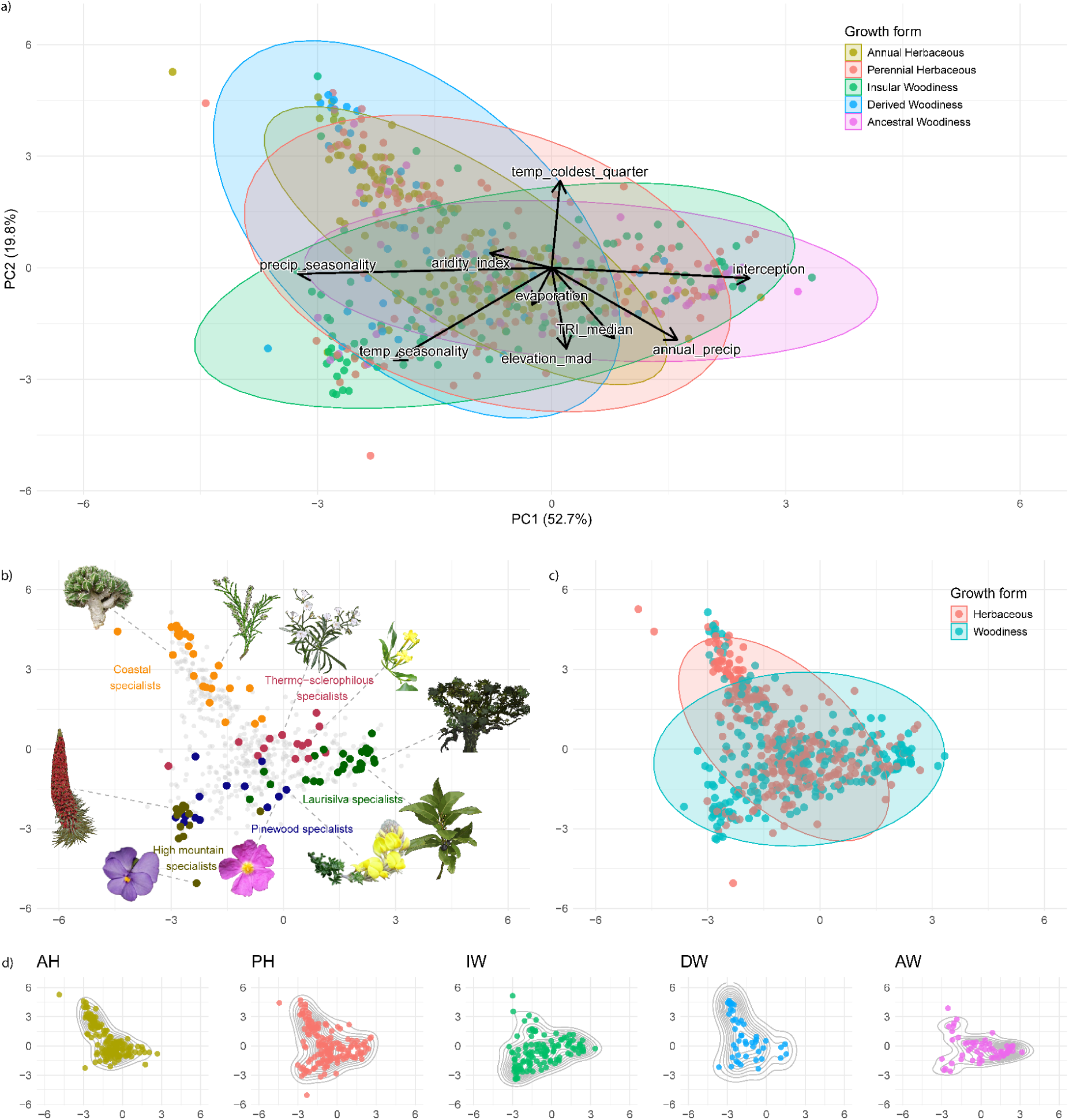
Environmental niche space of the growth forms of the Canary Island non-monocot angiosperm flora. a) Phylogenetic PCA of the different environmental variables labelled per growth form for the 548 species retained using the balanced sampling regime. Each point represents a species, shaded ellipses represent 95% confidence intervals around the points for each growth form, and the arrows represent the loadings of the phylo-PCA to highlight the contribution of each variable. b) Major Canarian vegetation zones for 99 species plotted onto the phylo-PCA, with a selection of specialists highlighted, using Perez (1999) and del Arco-Aguilar & Rodriguez (2018). Plant images from top-left, clockwise: *Euphorbia handiensis*, *Tamarix canariensis*, *Convolvulus floridus*, *Chrysojasminum odoratissimum*, *Ocotea [Mespilodaphne] foetens*, *Laurus novocanariensis*, *Adenocarpus foliolosus*, *Cistus symphytifolius*, *Viola cheiranthifolia*, *Echium wildpretii*. For the highlighted species labelled on the phylo-PCA, see Figure S7. c) Environmental niche space labelled in groups: herbaceous versus woody; and d) separated into five distinct growth forms: AH = Annual Herbaceous, PH = Perennial Herbaceous, IW = Insular Woody, DW = Derived Woody, AW = Ancestral Woody. Contour lines represent 2D kernel density estimation. For the sources of the plant images, see Table S4.

The Canary Islands host ∼1,375 angiosperm species, of which approximately a third are endemic to the archipelago (Canary Islands Biodiversity Database, 2024). This study focuses on angiosperms given their well-resolved taxonomic backbone (Sauquet & Magallón, 2018), their extensive representation in earlier evolutionary research (Hernández-Hernández & Wiens, 2020; Florencio et al., 2021), and the broad range of herbaceous (annual, perennial) and woody growth forms (insular woody, derived woody, ancestrally woody). We target all 1,160 non-monocot angiosperm species native to the Canary Islands.

#### 2.1.2. GBIF extraction & Locality data filtering

We developed an approach to extract, filter, and analyse publicly available occurrence and spatial data, whilst accounting for phylogeny, and to subsequently visualise environmental niche space. Details on our novel pipeline are covered in section 2.2 below. To obtain distribution data on the native non-monocot angiosperm species of the Canary Islands, we extracted occurrence data from the Global Biodiversity Information Facility (GBIF) using the ‘rgbif’ R package (Chamberlain et al., 2024; GBIF, 2025), following an updated version of Beierkuhnlein et al. (2021). For 186 plant species, no GBIF occurrences were found, and thus these were removed from the subsequent analyses, resulting in a total of 974 plant species for which at least one GBIF occurrence was found on the Canary Islands (Table S1). For a list of the changes made to Beierkunhlein et al. (2021), see Table S2

For more details on how we performed a subsequent filtering procedure to remove spatially erroneous occurrences associated with GBIF-mediated data (Maldonado et al., 2015; Zizka et al., 2020), we refer to Table S2, Supplementary Methods M1, and Table 1 in Supplementary Methods M1. An overview of the different functions used to filter the occurrence data can be found in Table SM1, including custom functions to filter occurrences reprojected from raster to vector data.

**Table 1.**
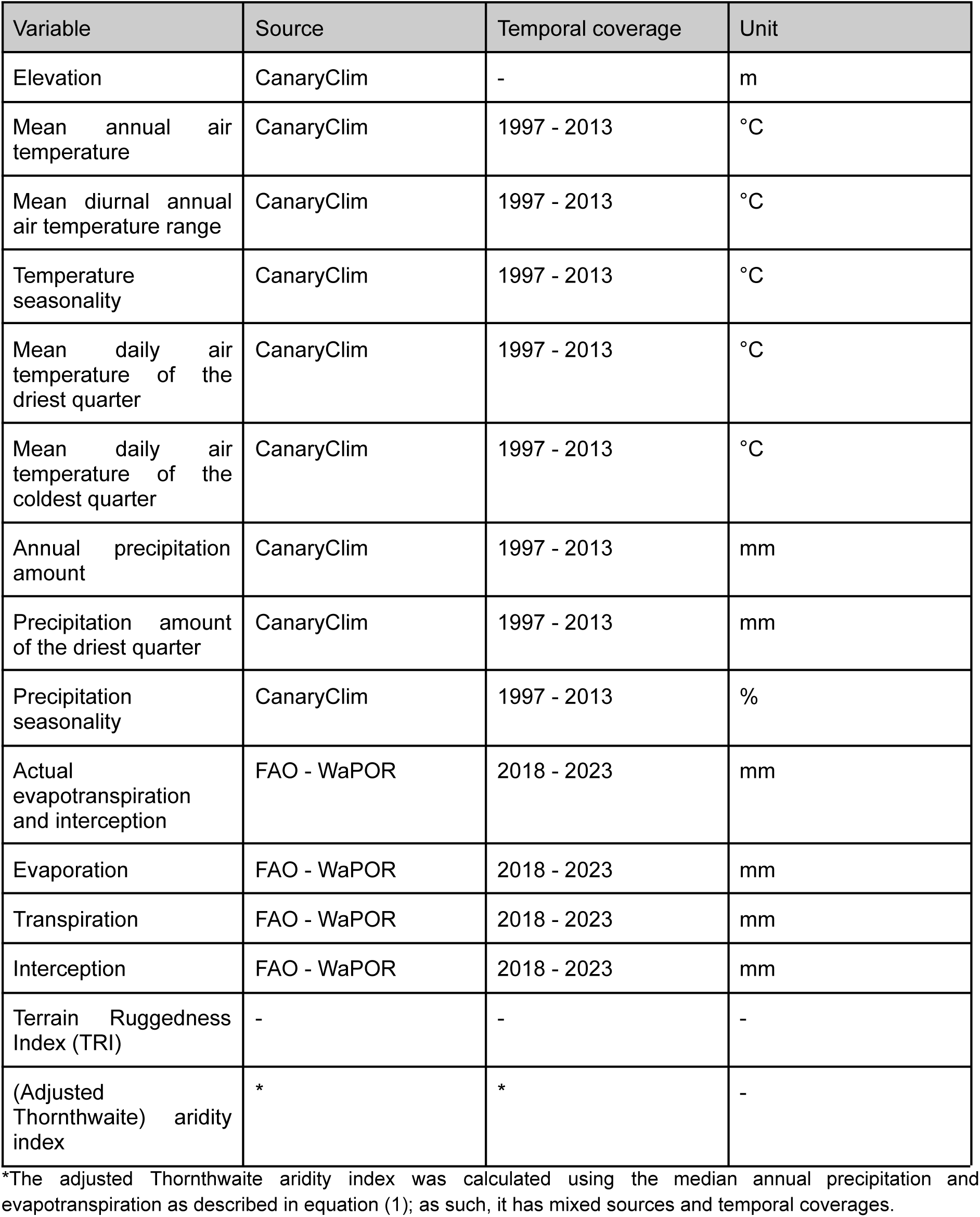
Bioclimatic and topographic variables used in this study.

#### 2.1.3. Sampling regimes

To find a balance between high species representation in terms of number of occurrence records and high coordinate certainty, we compared the results for three different uncertainty threshold resolutions: 1000, 500 and 100 meters, and minimum number of occurrences per species (after filtering): at least 3, 13 and 25 occurrences per species. These occurrence thresholds were based on van Proosdij et al. (2016), who suggested several thresholds to accurately model species distributions based on different geographic scenarios; fewer occurrences are needed when modelling species with small distributions (i.e., narrow endemics) than more widely distributed species.

These thresholds were combined into three sampling regimes: lenient, balanced, and strict. The lenient sampling regime has a minimum of three occurrences per species with a maximum uncertainty limit of 1000 meters; the balanced sampling regime has at least 13 occurrences with a maximum uncertainty of 500 meters; and the strict sampling strategy has at least 25 occurrences and a maximum uncertainty of 100 meters.

### 2.2 Environmental dataset compilation

#### 2.2.1. Variable extraction

We extracted 15 environmental variables for the (filtered) occurrences for each species from various environmental (raster) layers from multiple sources. An overview of these variables and their sources is presented in Table 1. All environmental layers had a resolution of 100 meters (∼3-4 arc-seconds) and were assigned the same coordinate reference system. See the ‘Code Availability’ section for the source files. For more information on how these environmental variables were selected and from which source they were extracted, see Table 1 and Supplemental Methods M1.

#### 2.2.2. Variable aggregation

After extracting the 15 environmental variables for each species, we aggregated the occurrence-level data to the species level. To summarise the information from occurrences, we used the median value for most environmental variables (Table 1), because of its robustness against outlying values (Müller, 2000). For elevation, besides the median, we also used the median absolute deviation (MAD) to investigate the relationship between the elevational range at which a species occurs and its growth form.

#### 2.2.3. Growth form determination

For each species, we scored the growth form in one of five categories: annual herbaceous (AH), perennial herbaceous (PH), insular woodiness (IW), derived woodiness (DW), and ancestral woodiness (AW), following Hooft van Huysduynen et al. (2021). Details on how the growth form types were scored are provided in Supplemental Methods M1. See Table S1 for the growth form scoring for all the species included in this study.

### 2.3 Environmental data analysis

#### 2.3.1 Correlation Analysis

We performed Kendall’s correlation tests to remove strongly correlated environmental variables (τ > 0.7; Figure S1), and ran the analyses using the balanced sampling approach. To allow for comparison between results, we used the same subset of nine variables that were retained for subsequent analyses using the strict and lenient approaches.

#### 2.3.2. Phylo-PCA & Phylo-ANOVA

To visualise the major axes of variation across the environmental variables, while accounting for the shared evolutionary history among taxa, we performed phylogenetic principal component analysis (hereafter ‘phylo-PCA’) (Revell 2009, 2024) (see Supplemental Methods M1). To test for differences in the environmental variables across the five different growth forms, while accounting for their shared evolutionary history, we performed a phylogenetic Analysis of Variance (phylo-ANOVA; Figure 3) analysis (see Supplemental Methods M1).

**Figure 3.**
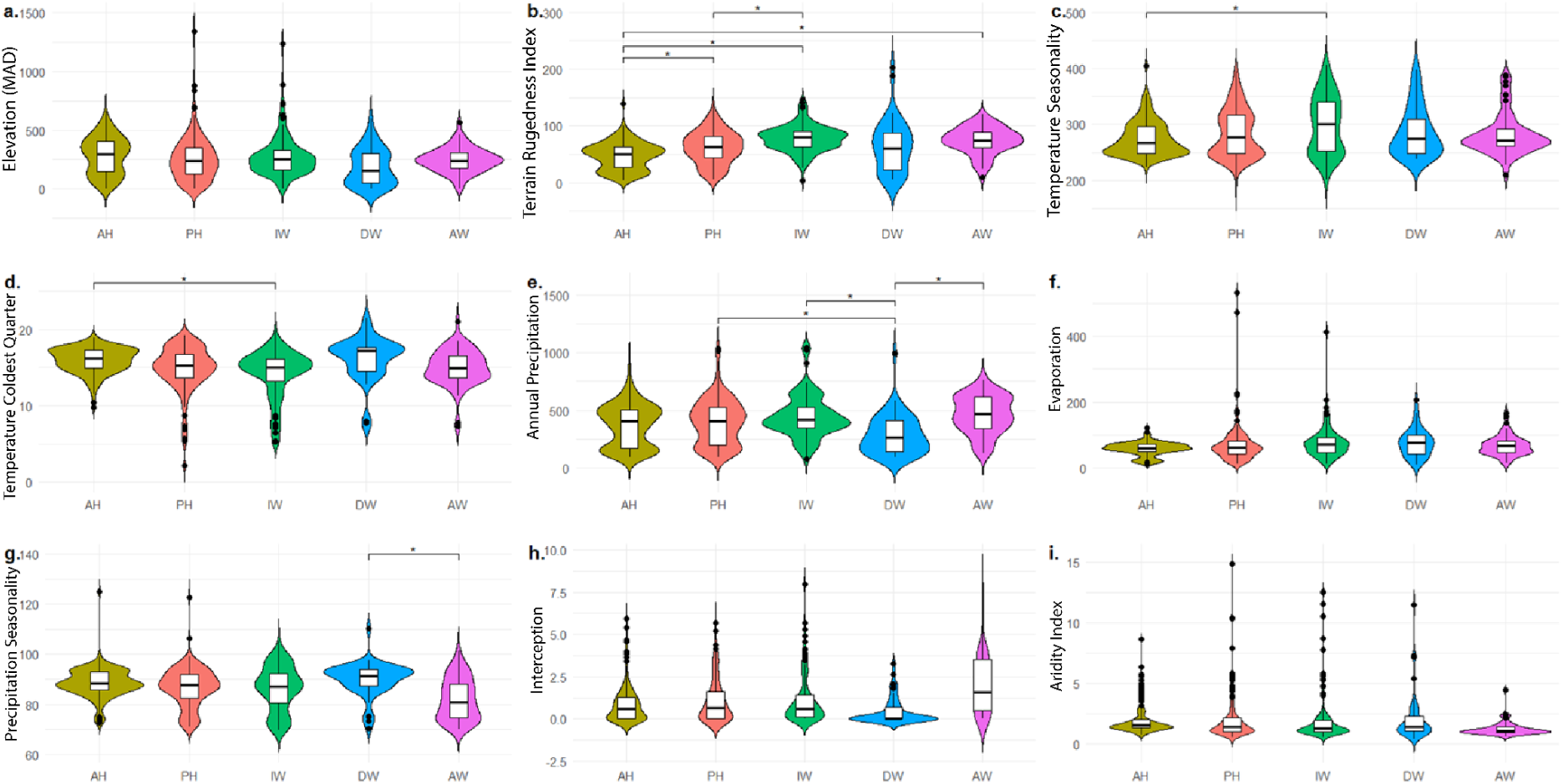
Results of the phylo-ANOVA. Boxplots show the distribution of different environmental variables across each growth form: a) Elevation (MAD), b) Terrain ruggedness index, c) Temperature seasonality (°C*100), d) Temperature in the coldest quarter (°C), e) Annual precipitation (mm), f) Evaporation (mm), g) Precipitation seasonality (%), h) Square root of the interception (mm), and i) Adjusted Thornthwaite aridity index. The violin plots show the distribution of each variable per growth form and the median value for each environmental variable for each growth form. AH = annual herbaceousness, PH = perennial herbaceousness, IW = insular woodiness, DW = derived woodiness, AW = ancestral woodiness. Significant group differences (Holm-adjusted p-value < 0.05; phylo-ANOVA) are indicated with *. The square root of the interception is used due to the spread of the raw data.

#### 2.3.3. Rugosity (La Palma case study)

To further explore the differences between growth forms found by the phylo-ANOVA, we plotted the base terrain ruggedness index (TRI) raster layer against the proportion of the occurrences of each growth form, relative to the total occurrences, per grid cell for the island of La Palma (Figure 4). We used La Palma as a case study due to the clarity of the valleys and ravines on the rugosity raster layer compared to the other islands. We used the ‘geom_sf’ package in ggplot2, where each grid cell corresponds to ∼300m^2^ (Wickham, 2016). Visualising the proportion of each growth form per grid cell, relative to the total occurrences within the grid cell, highlights regions where one growth form predominates. Visualising the raw raster file also provides context for how gradients observed over topographically complex regions relate to the distribution of each growth form. For the same analyses repeated across the other Canarian islands (except Lanzarote and Fuerteventura due to the lack of variation in topographic complexity), see Figures S2-5.

**Figure 4.**
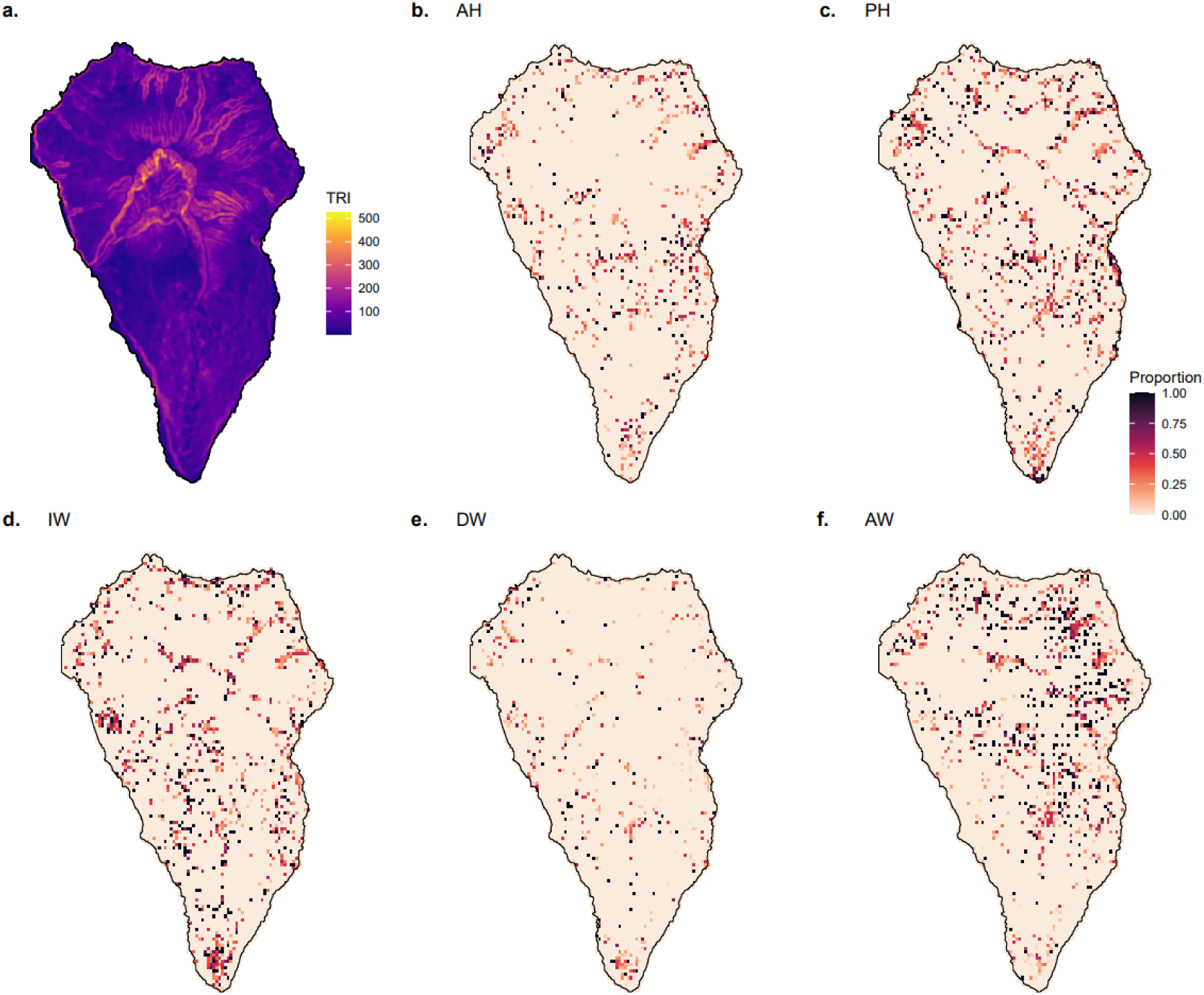
Rugosity in La Palma Island and the distribution of different growth forms, using the proportion of each growth form relative to the total occurrences per grid cell. a) Raster of terrain ruggedness index, b) AH = annual herbaceousness, c) PH = perennial herbaceousness, d) IW = insular woodiness, e) DW = derived woodiness, and f) AW = ancestral woodiness. Peach grid squares can refer to zero proportion of a growth form or a grid cell with no data. For the distribution of the growth forms on the other Canary Islands, see Figures S2-5.

## 3. Results

### 3.1 Locality Data Extraction

Of our 1,160 target species, we obtained growth form data for 1,155 species, and 974 of these had locality data in GBIF (Table 2, Table S1). Table 2 shows the number of species retained after filtering for each of the three sampling regimes across all five growth forms.

**Table 2.** The number of species targeted in this study, with locality data in GBIF, and included in each sampling approach. (lenient, balanced, and strict), with the percentage of species with locality data remaining after filtering for each sampling regime.

| Growth Form | Total in target list | Total with GBIF data | Lenient approach | % after filtering (lenient) | Balanced approach | % after filtering (balanced) | Strict approach | % after filtering (strict) |
| --- | --- | --- | --- | --- | --- | --- | --- | --- |
| Annual Herbaceous | 371 | 297 | 231 | 77.8 | 153 | 51.5 | 91 | 30.6 |
| Perennial Herbaceous | 320 | 263 | 222 | 84.4 | 161 | 61.2 | 126 | 47.9 |
| Ancestral Woodiness | 126 | 105 | 89 | 84.8 | 63 | 60.0 | 55 | 52.4 |
| Derived Woodiness | 97 | 85 | 67 | 78.8 | 47 | 55.3 | 33 | 38.8 |
| Insular Woodiness | 241 | 224 | 194 | 86.6 | 124 | 55.4 | 91 | 40.6 |
| Total | 1155 | 974 | 803 | 82.4 | 548 | 56.3 | 396 | 40.7 |

The percentage of species remaining from the total number of species for which GBIF locality data were extracted is also reported. As expected, the lenient sampling regime has the most species retained after filtering, with 803 of the 974 species retained (82%); the strict sampling regime has the fewest species retained, at 396 (41%); and the balanced sampling regime retains 548 species (56%).

The results and figures in the following sections are shown for the balanced sampling regime, unless stated otherwise (for results from lenient and strict sampling regimes, see Figures S8 & S9). A minimum number of 13 occurrences (as in the balanced regime) implies both narrow endemic species as well as more widespread species are included, following van Proosdij et al. (2016), and an uncertainty limit of 500 m balances spatially accurate distributions with high species representation. However, we acknowledge that given the high environmental variability within the Canary Islands, 500 m resolution may be too coarse to capture the true climatic conditions of species.

### 3.2 Environmental Niche Space (Phylo-PCA)

To investigate the environmental variables associated with different plant growth forms in the Canary Islands and address our first and second hypotheses, we used a phylo-PCA to map the environmental niche space of five growth forms (Figure 2, Table S3). We find a broad overlap between the growth forms’ environmental niche spaces (confidence ellipses), aggregating in the centre of island environmental niche space (Figure 2a). However, there are some distinctions in the direction of environmental niche space occupied by each of the growth forms (Figure 2a), suggesting that the environmental variables have differing importance for each growth form. For example, IW species are placed mostly along a climatic seasonality axis, linked to both precipitation and temperature (PC1 in Figure 2), and to a minor degree, rugosity (TRI_median). The IW environmental niche space shows strong convergence with the AW niche, though the latter is less linked to increased seasonality and more related to increased interception, i.e. the precipitation caught by vegetation that evaporates before reaching the ground. In contrast, the niche space of herbaceous species is mostly dispersed along the temperature-seasonality-elevation-precipitation axis (PC2 in Figure 2a), with precipitation seasonality (PC1 in Figure 2) more important in PH than AH. Unexpectedly, DW niche space shows greater congruence with the herbaceous growth forms than with the other woody growth forms; a finding likely attributed to DW species occurring more prevalently in coastal regions than other woody growth forms (Figure 2a, 2b). For how each variable contributes to the principal component axes, see Figure S6.

To assess the validity of the extracted environmental variables (Table S3), we compared our recovered environmental niche space to the main vegetation zones present in the Canaries following Perez (1999) and del Arco-Aguilar & Rodríguez (2018): (dry) coastal region, high-altitude region, laurisilva forest, pine forest region, and the thermo-sclerophyllous forests. We highlight ∼100 species that are specialists of or are typical of a single vegetation zone in niche space (Figure 2b) to show the phylo-PCA largely clusters species into coherent regions (Perez, 1999; del Arco-Aguilar & Rodríguez, 2018): coastal species cluster in arid regions of low precipitation and low rugosity, given by smaller PC1 values and larger PC2 values (Figure 2a, S2, Table S3), and laurisilva specialists are found in areas of high interception (indicative of closed canopy cover) and low seasonality, given by more positive PC1 values (Figures 2a, 2b, S2, Table S3).

To further explore broad differences in niche space and test our first hypothesis, we aggregated our growth form data into binary woody and herbaceous growth forms (Figure 2c). As with Figure 2a, overlap in niche space is observed centrally, but the niche space can be separated along the temperature-seasonality-elevation-precipitation axis (PC2), likely resulting from the dominance of herbaceous species in coastal regions.

When subdividing our phylo-PCA into five plots by growth form from AH towards AW, a slight step-wise shift away from a predominance in arid and aseasonal niche space typical of AH species becomes visible (top-left of niche space in Figure 2d, given by high PC2 values). Progressively, niche space gently shifts towards rugose, seasonal regions typical of IW species (bottom-left in Figure 2d, given by low PC1 values), or aseasonality and high interception (bottom-right in Figure 2d, given by high PC1 values) in AW species, whilst DW species are widely distributed across niche space.

The results in Figure 2 (using the balanced sampling regime) are largely congruent with those using other sampling regimes, suggesting 500 m coordinate uncertainty may not be too coarse to capture climatic preferences of species. Across sampling regimes, overlapping clusters are observed centrally, with differences between niche space found largely in the directionality of the confidence ellipses. In the strict sampling regime, the directionality of the niche space for each growth form was further pronounced, with greater convergence between AH and PH niche space, and between IW and AW niche space (Figure S8). DW niche space remains widespread throughout niche space, but with increased overlap with the AH niche. For the lenient sampling regime, there is greater overlap between the growth forms, with little distinction between the herbaceous and woody niche in niche space (Figure S9), but the relative importance of each variable on each growth form remains similar.

### 3.3 Pairwise comparisons (phylo-ANOVA)

To statistically analyse the differences between the localities where the growth forms occur, in terms of the environmental variables, and test our third hypothesis, we used a phylo-ANOVA performing pairwise comparisons between each growth form for each environmental variable (Figure 3). We find significant differences between at least two growth forms in five of the nine variables analysed. Rugosity has the most pairwise significant differences between growth forms, with significance found in four pairwise tests (Figure 3b). Generally, AH species have the lowest rugosity, with rugosity being higher in woody growth forms than herbaceous ones, and IW species having the highest observed median rugosity. Despite both temperature and precipitation seasonality shaping environmental space (Figure 2a), there is only one significant pairwise difference for each of these variables (Figure 3c, g). IW species have significantly higher temperature seasonality than AH species, with the highest median temperature seasonality across all growth forms (Figure 3c). AW species have significantly lower precipitation seasonality than DW species, the former having the lowest observed median precipitation seasonality and the latter the highest (Figure 3g). Three significant pairwise differences are found for annual precipitation (Figure 3e), between DW and PH, DW and IW, and DW and AW. IW species are also found in significantly colder regions (in the coldest quarter) than AH species (Figure 3d).

### 3.4. Rugosity (La Palma case study)

As the phylo-ANOVA revealed rugosity can best separate the five growth forms (Figure 3b), we explored the differences between growth forms along the rugosity variable further (Figure 4). By plotting the base TRI raster alongside the proportion of each growth form per grid cell, we can visualise the relationship between rugosity and areas where each growth form is particularly prevalent. We find that high proportions of IW species occurrence are found in high rugosity areas, such as around the central Caldera de Taburiente and the ravines and narrow valleys with steep flanks scattered in the northern and central part of the island (Figure 4d). The high proportion of PH species in highly rugose areas (Figure 4c), as well as across the lower rugosity southern regions of La Palma, reflects the wide range of TRI values obtained for this growth form (Figure 3b). Figure 4 also highlights the predominance of AW species within the (rugose) laurisilva forest of La Palma (Figure 4f), and the preference of AH species for drier, coastal areas of lower rugosity (Figure 4b), in agreement with the findings of the phylo-PCA (Figure 2). Rugosity, therefore, emerges as an important variable for the distribution of growth forms in the Canary Islands.

## 4. Discussion

Here, we present a comprehensive analysis of the associations between phylogenetically inferred growth forms and environmental variables for the Canary Islands’ (Figure 1) non-monocot angiosperm community, comprising 1,160 native species (Figure 2). Using a novel bioinformatics pipeline, we build on existing locality data-filtering methods by incorporating functions to remove occurrences on islands to which a given species is not native or to which occurrence records are reprojected from raster data into vector data, and allowing for selecting pre-defined thresholds of coordinate uncertainty and number of occurrences. After extracting and filtering GBIF occurrence data for 974 species of the non-monocot flora (84%), we extracted and aggregated environmental data for each species (Table S3), and visualised environmental niche space for the 548 species that passed data filtering using a phylo-PCA (Figure 2). For the final steps of our approach, we used phylo-ANOVAs to statistically analyse the pairwise differences between the environmental variables for each growth form to establish which variables are important for segregating the niche space of these growth forms (Figures 2, 3).

We highlighted 99 species on the phylo-PCA that occur in a single major Canarian vegetation zone (Figure 2b), following Perez (1999) and del Arco-Aguilar & Rodriguez (2018), mapping the species’ names, which suggests our pipeline produces results consistent with known habitat occurrences (Figure S7). For example, known specialists of high montane systems such as *Echium wildpretii* and *Viola cheiranthifolia* are recovered well within regions of niche space linked to high elevation and seasonality, given by small PC1 and PC2 values (Figures 2b, S2, Table S3; Rodríguez-Rodríguez et al., 2019; Graham et al., 2021). The iconic cactiform *Euphorbia* species typical of the xerophytic lowland communities (“cardonal-tabaibal”), such as *E. canariensis* and *E. handiensis* (Coello et al., 2024), are mapped in high aridity, and low elevation and rugosity areas of niche space. Endemic species of the laurisilva subtropical forest, such as *Laurus novocanariensis* and *Ocotea foetens*, occur in regions of niche space dominated by high precipitation and interception (precipitation caught by vegetation that evaporates before reaching the ground), with low seasonality (Figure 2a, Table S3; Betzin et al., 2016). Our niche space also resembles dos Santos et al. (2022), who find a triangular shape with coastal regions, high montane systems, and the laurisilva forests on the corners.

Though there are several existing pipelines to clean and analyse occurrence data (e.g., Zizka et al., 2019), and subsequently extract environmental variables from spatial data (van den Hoogen et al., 2021; Velazco et al., 2022; Coca-de-la-Iglesia et al., 2023; Jemeļjanova et al., 2024), our approach goes beyond these first steps by incorporating existing phylogenetically-corrected visualisation and statistical methods. This means our approach allows us to visualise environmental niche space for any predefined set of categorical traits (e.g., growth form, but taxonomy/clade can be easily implemented), rather than only exploring the extracted environmental data in a species distribution modelling context to predict occurrence as implemented in existing pipelines (van den Hoogen et al., 2021; Velazco et al., 2022). Our method also allows for greater precision in delimiting the native range of a species than in existing methods based on the geographical codes of the Taxonomic Databases Working Group (TDWG; Brummitt, 2001). For example, by using island-level native status across the Canary Islands from published floras (Beierkuhnlein et al., 2021; Canary Islands Biodiversity Database, 2024), we can more accurately map the native range of a species across the archipelago than methods that use TDWG codes that would treat the Canary Islands as a single region. Finally, as we use floras as input in our approach, we provide an alternative method for accounting for taxonomy and synonymy than the previously existing ‘scrubr’ R package (Chamberlain, 2025).

### Substantial overlap in environmental niche space for growth form types in the Canary Islands flora

We hypothesised that the marked differences in life span, woodiness, and evolutionary history across the five growth forms should lead to pronounced differences in environmental niche space. However, we found considerable niche overlap across the five growth forms centrally in niche space (Figure 2), e.g. as can be seen by the confidence ellipses (Figure 2a). For example, annual and perennial herbaceous growth forms (AH, PH) show marked overlap, despite differences in life span, and derived woody species (that acquired their woodiness before they colonised the islands; DW) overlap considerably with AH species (Figure 2a), likely explained by the predominance of both growth forms in coastal areas (Figure 2a, b). Nevertheless, the niche space of the growth forms does show some distinctions, providing mixed support for our first hypothesis (that there will be detectable differences in niche space between herbaceous and woody species, and additional differences between the five growth forms). Interestingly, in support of the second hypothesis (that the insular woody (IW) niche will deviate most from those of herbaceous growth forms), IW species exhibit distinctions from the herbaceous growth forms (Figure 2a), and show strong convergence to occupy similar environmental space as ancestrally woody (AW) species, although differing in the effect of seasonality. This marked convergence in niche space between IW and AW, despite woodiness evolving differently across evolutionary history (i.e., from a herbaceous ancestor in IW species), hints at the existence of a ‘woody niche’ in niche space (Lens et al., 2013a). However, following this line of thought, we would expect the woody niche to be more distinct as postulated in our first hypothesis (Figure 2c). We do find some support for this when the niches of the three grouped woody growth forms are compared with the two herbaceous growth forms, but the woody-herbaceous niche overlap remains considerable (Figure 2c). These findings are also robust under different sampling regimes (Figures S8, S9).

Unexpectedly, and contrasting previous studies and our third hypothesis (that variables related to aridity and seasonality will impact the niche of IW species; Carlquist, 1974; Zizka et al., 2022), IW species are found in areas of higher climatic seasonality (Figures 2a, d), given by smaller PC1 values (Figure 2a, S2, Table S3). Most variation in the niche space of IW species is explained by the precipitation-seasonality-interception axis (PC1 in Figure 2, Figure S6), and IW species exhibit the highest overall median temperature seasonality (Figure 3c), as well as the lowest overall median temperature in the coldest quarter (Figure 3d). This suggests greater tolerance to seasonally lower temperatures than herbaceous species, contrasting global studies (Klimeš et al., 2022). However, none of the IW species analysed occur in areas where frost occurs, as none of them grow in regions where the temperature in the coldest quarter reaches the frost point (Figure 3d). An aseasonal climate, defined by Carlquist (1974) as growth uninterrupted by frost, is therefore still characteristic for the IW Canary Islands species, and could still serve as a potential driver favouring woodiness, despite seasonality being important in shaping the IW environmental niche space (Figures 2a, 3g). Another perhaps more unexpected finding of our study is that the IW niche is not defined by parameters indicating increased drought, contrary to previous studies (Lens et al., 2013b; Doria et al., 2018; Hooft van Huysduynen et al., 2021; Zizka et al., 2022). This does not exclude the possibility, however, that drought did play an important role in promoting the evolution of woodiness on the Canary Islands for at least some lineages (e.g., Doria et al., 2018) and serves as an important global woodiness driver on continents.

Upon comparing the other growth forms used in this study to previous studies, we recover several established relationships. For instance, AH species favour arid regions of low precipitation and elevation, and consistently high temperatures (Figures 2a, d, 3c, d, Table S3), consistent with previous analyses (Irl et al., 2020; Poppenwimer et al., 2023), and when previously assessed as therophytes (Taylor et al., 2023). Also, PH species, previously assessed as hemicryptophytes or subshrubs, show similar correlates as AH species but with greater importance of increasing precipitation and elevation, and lower temperatures (Irl et al., 2020; Taylor et al., 2023). This relationship is also recovered here (Figure 2a), as given by the PH niche space being shaped more by PC2, the temperature-seasonality-elevation-precipitation axis, than AH species (Figure S6). Likewise, high precipitation and low climatic seasonality (both precipitation and temperature) are correlated with the distribution of AW species (Figures 2, 3c, e, g), explaining nearly all variation across PC1 (Figure 2), in agreement with previous studies who classify AW species as phanerophytes and trees, respectively (Poppenwimer et al., 2023; Taylor et al., 2023).

An important trait that best shapes plant growth form in the Canaries is topographic complexity, included in our approach as rugosity (TRI variable). IW species are those that are found to occur generally in areas with the highest rugosity, significantly higher than both herbaceous growth forms and significantly higher in AW than AH species (Figure 3b). Additionally, PH species are found in more topographically complex areas than AH species (Figure 3b). These findings demonstrate that woody species, and to a lesser extent perennials, are abundant in more topographically complex regions, which could be explained by two factors. First, the current distributions of woody species in highly rugose areas may be relict populations, reflecting grazing pressure from large mammal herbivores introduced to the archipelago over the last ∼2,000 years (Castilla-Beltrán et al., 2021). Alternatively, over longer evolutionary timescales, topographically complex regions may create microclimates and/or micro-habitats, which could spur speciation events due to limited seed dispersal, suppressed gene flow between isolated populations, and increased climatic diversity along steep elevational gradients (Vitales et al., 2014; Chang et al., 2023; Graham et al., 2025). Therefore, the recovered relationship between (insular) woodiness and rugosity may stem from a combination of a lack of herbivory in these potentially relict populations affecting extant distribution patterns, and the presence of IW species in areas of increased opportunity for allopatric speciation. As most IW species on the Canaries belong to evolutionary radiations (Carine et al., 2010; Lens et al., 2013b), other factors that promote diversification within these lineages may allow for rapid diversification along the steep elevational gradient found in highly rugose areas, thereby creating the correlation between rugosity and IW.

In conclusion, based on our approach, spanning the extraction of specimen records to phylogenetically-inferred comparison and visualisation of environmental niche space, we find substantial overlap across the niches of five plant growth forms on the Canaries, with significant differences in the environmental correlates of various woody and herbaceous growth forms across the Canary Islands. Climatic seasonality (both temperature and precipitation) and topographic complexity emerge as key distinguishing factors in driving differences in environmental niche space. Although the environmental niche space overlap across the growth forms is unexpectedly high given the differences in life span and evolutionary history, our results for AH, PH and AW species confirm previous results based on global analyses. Surprisingly, for IW species, the environmental niche space is shaped by seasonal colder (but frost-free) temperatures without any correlation to drought-related parameters. Our results also highlight the importance of separating species into growth forms congruent with their evolutionary history (e.g. ancestral woodiness versus insular woodiness), as derived woody island species show greater similarity in their environmental niche to herbaceous species than to other woody growth forms, with the former primarily linked to temperature, elevation, and precipitation (or lack thereof). Overall, we hope our approach, which can be applied to any categorical trait of interest, will promote phylogenetically-inferred macroecological studies across the world, thereby contributing to how traits have shaped environmental niche space for any plant-animal-fungi lineage.

## 5. Author Contributions

Conceptualisation: L.J., R.F.A.B., L.V., F.L.; Environmental variable selection: L.J., R.F.A.B., J.C.-C., L.V., F.L.; Analyses: L.J., R.F.A.B.; Writing first draft: L.J., R.F.A.B., L.V., F.L.; Supervision: R.F.A.B., L.V., F.L. All co-authors contributed to editing and reviewing the manuscript.

## 6. Code Availability

The code and data required to recreate all analyses can be found in the supporting GitHub repository at https://github.com/ljn261/canary_islands_project.

## 6. Supporting information

**Figure S1.**
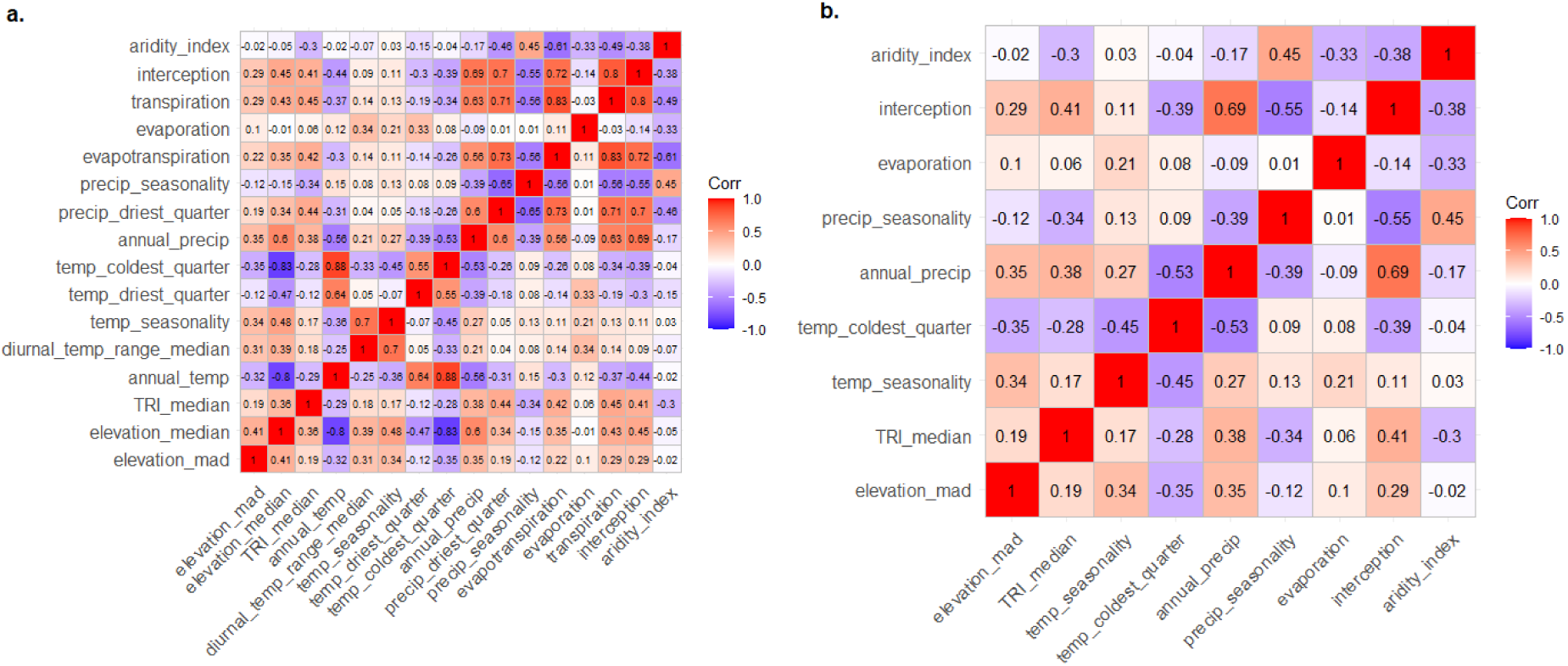
Correlation matrix using Kendall’s tau statistic with all variables. (a) and after variable selection (b).

**Figure S2.**
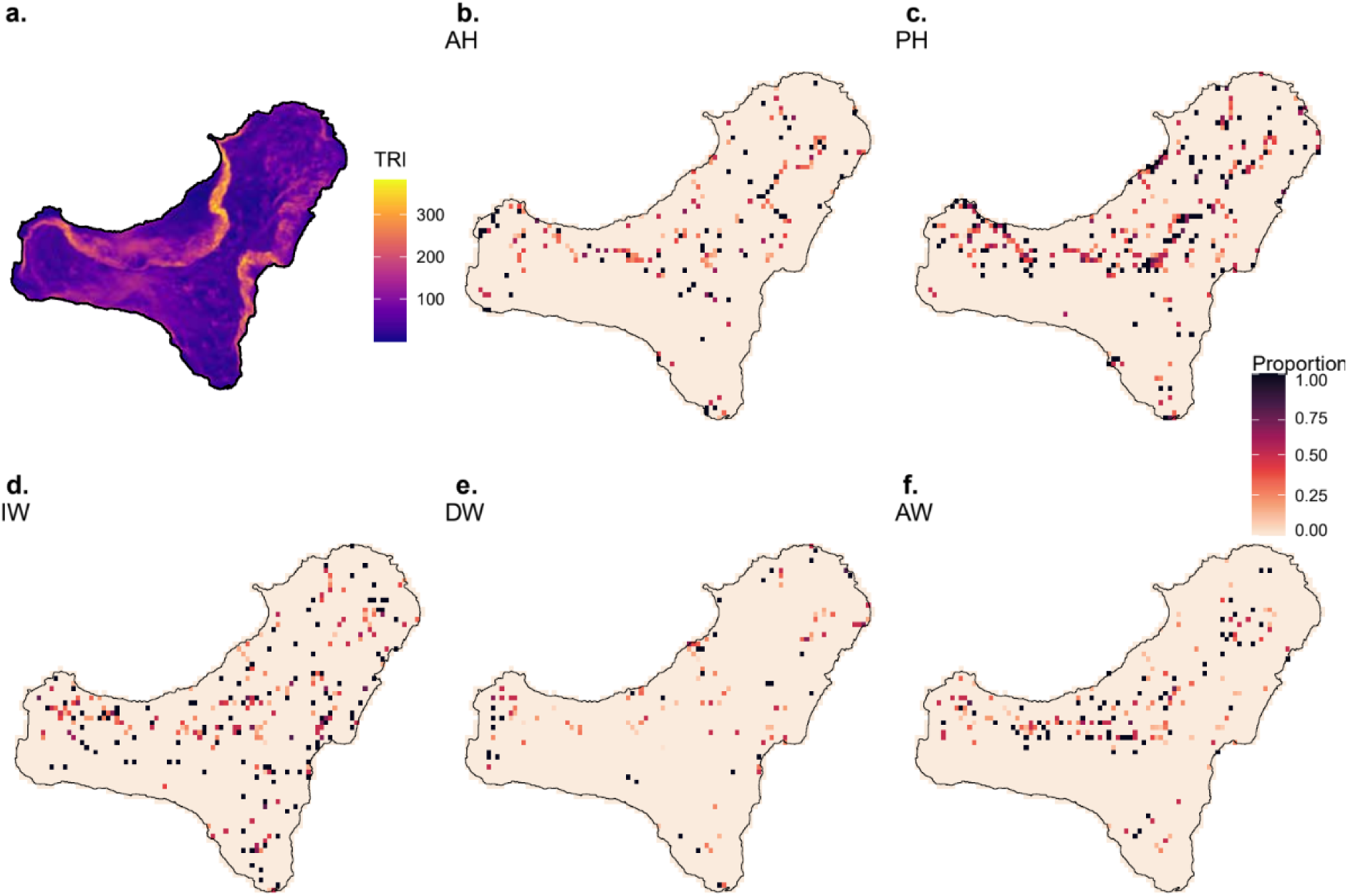
Rugosity (a) in El Hierro Island and the distribution of different growth forms (b-f), using the proportion of each growth form relative to the total occurrences per grid cell.

**Figure S3.**
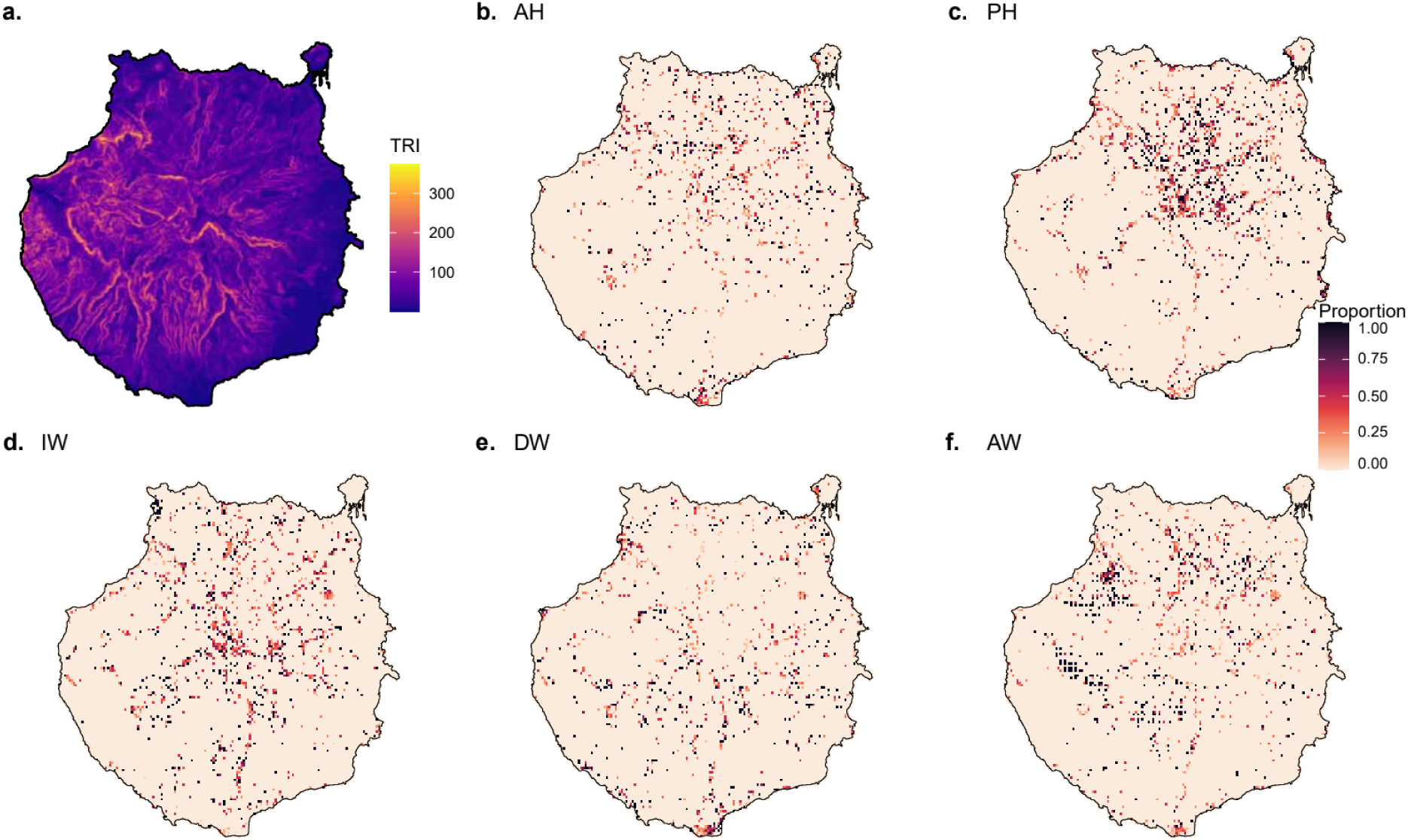
Rugosity (a) in Gran Canaria Island and the distribution of different growth forms (b-f), using the proportion of each growth form relative to the total occurrences per grid cell.

**Figure S4.**
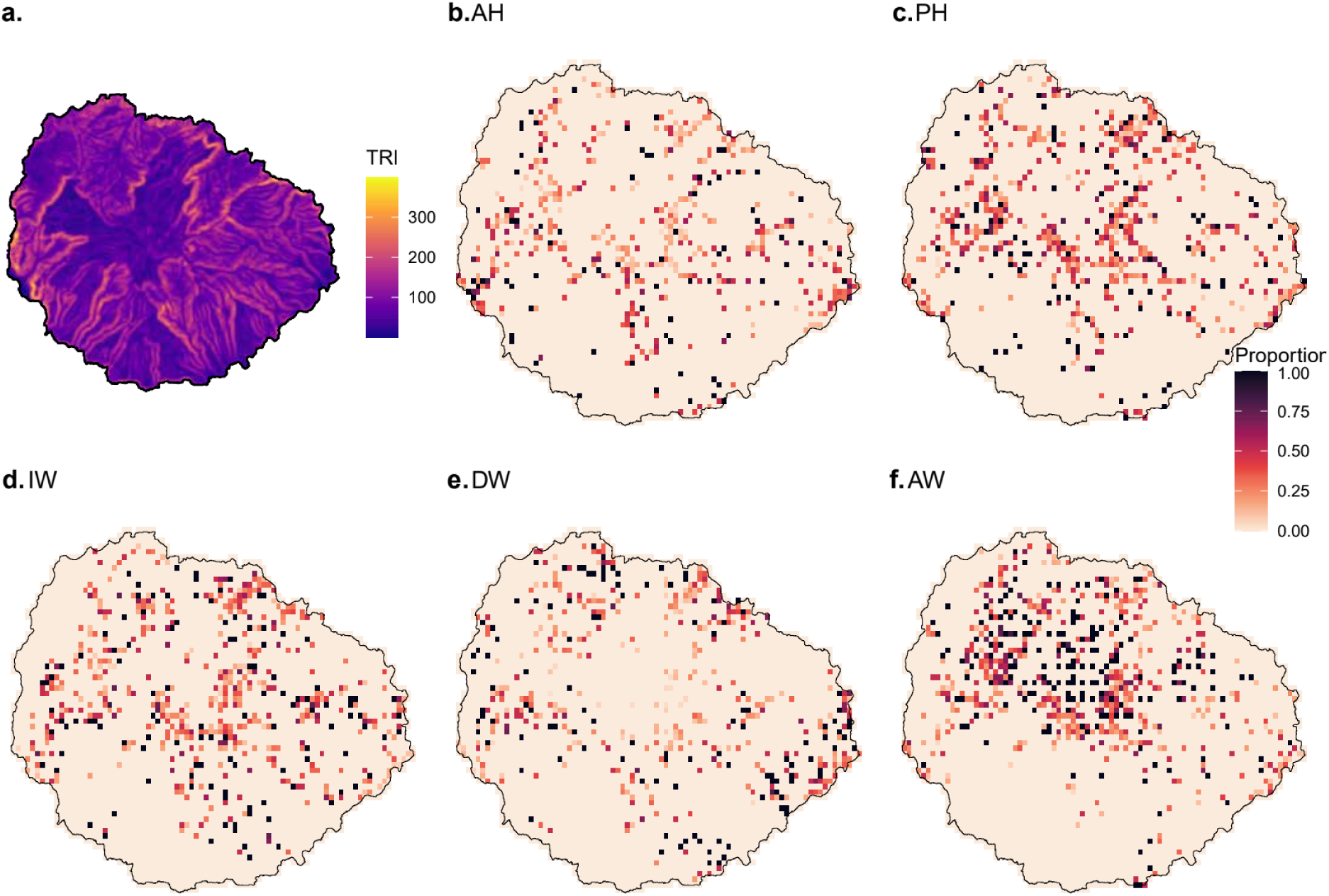
Rugosity (a) in La Gomera Island and the distribution of different growth forms (b-f), using the proportion of each growth form relative to the total occurrences per grid cell.

**Figure S5.**
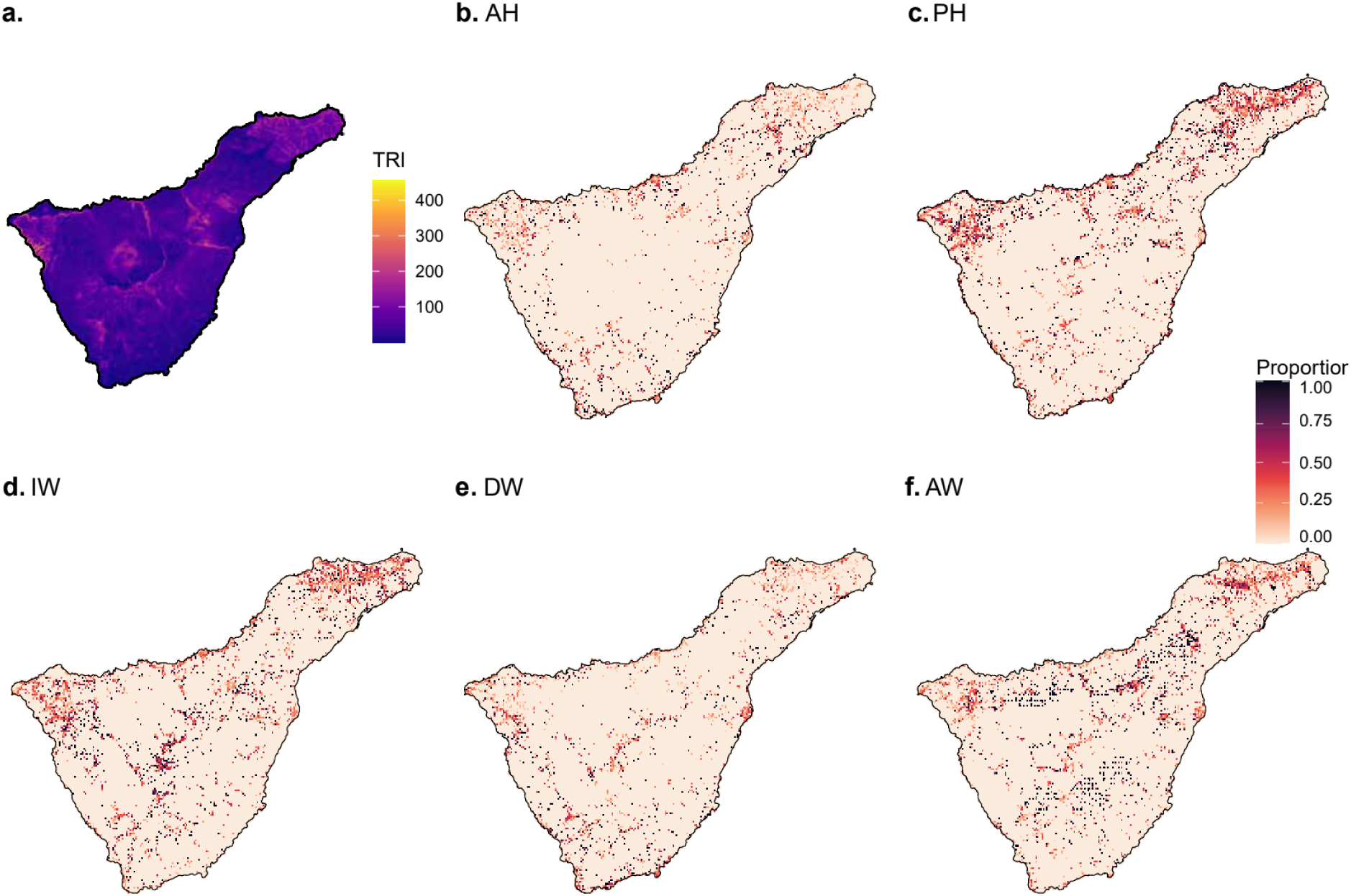
Rugosity (a) in Tenerife Island and the distribution of different growth forms (b-f), using the proportion of each growth form relative to the total occurrences per grid cell.

**Figure S6.**
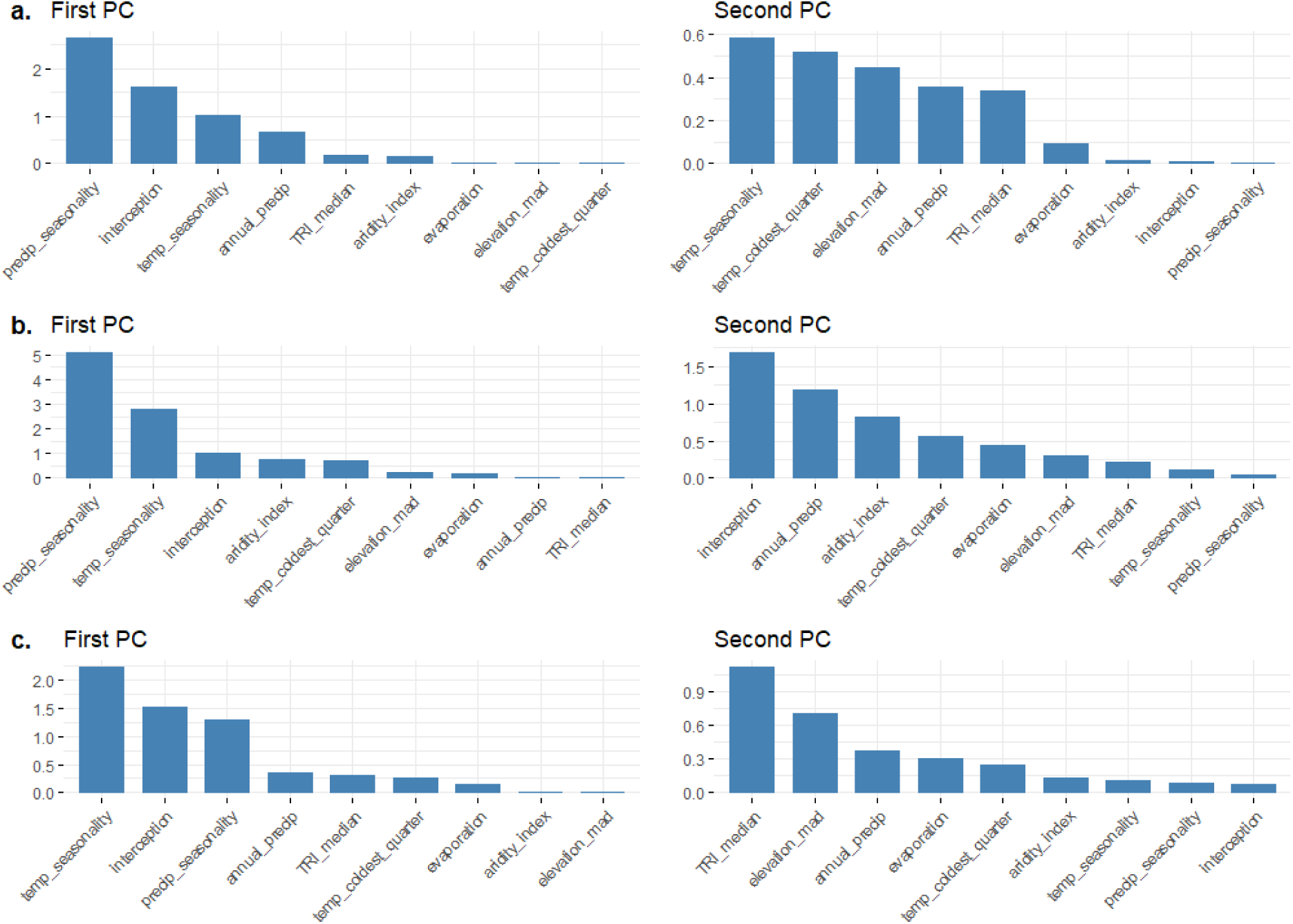
Squared cosine contribution to principal component axes, showing quality of representation in PCA run with balanced (a), lenient (b) and strict (c) sampling approach data

**Figure S7.**
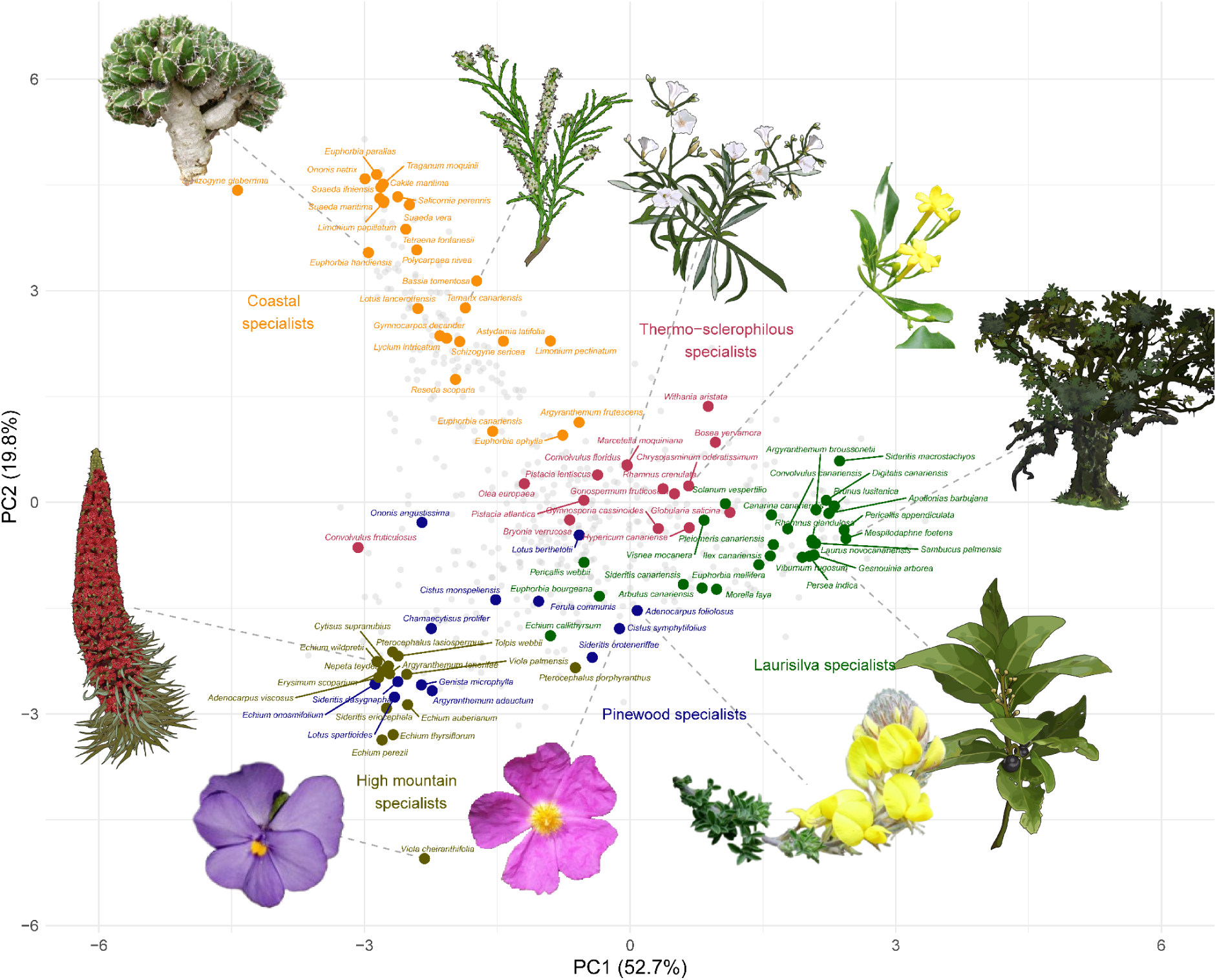
The major Canarian vegetation zones for 99 species plotted onto the phylo-PCA from Figure 2b, with the species labelled on the phylo-PCA. All species habitat information taken from Perez (1999) and del Arco-Aguilar & Rodriguez (2018).

**Figure S8.**
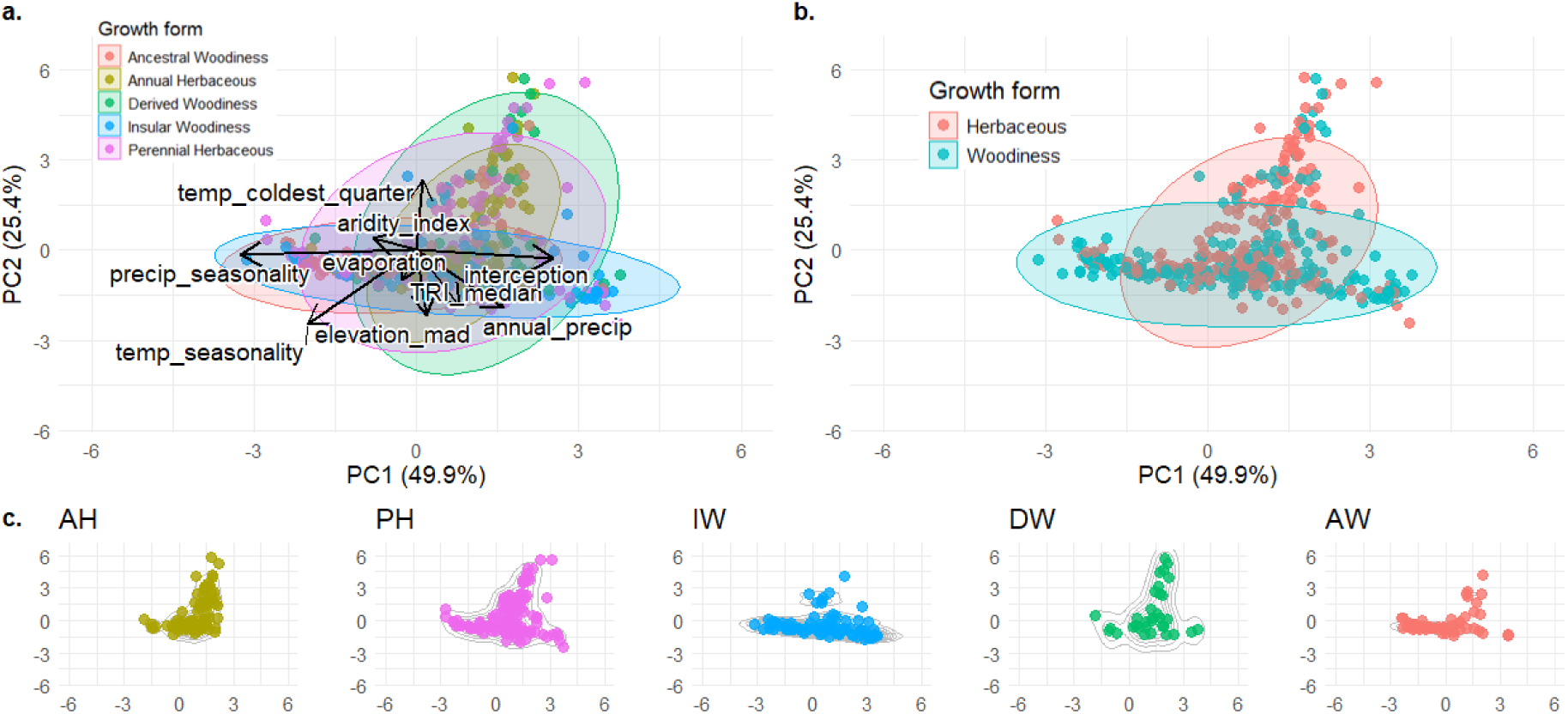
PCA results of the strict sampling approach. Formatted in the same way as Figure 2.

**Figure S9.**
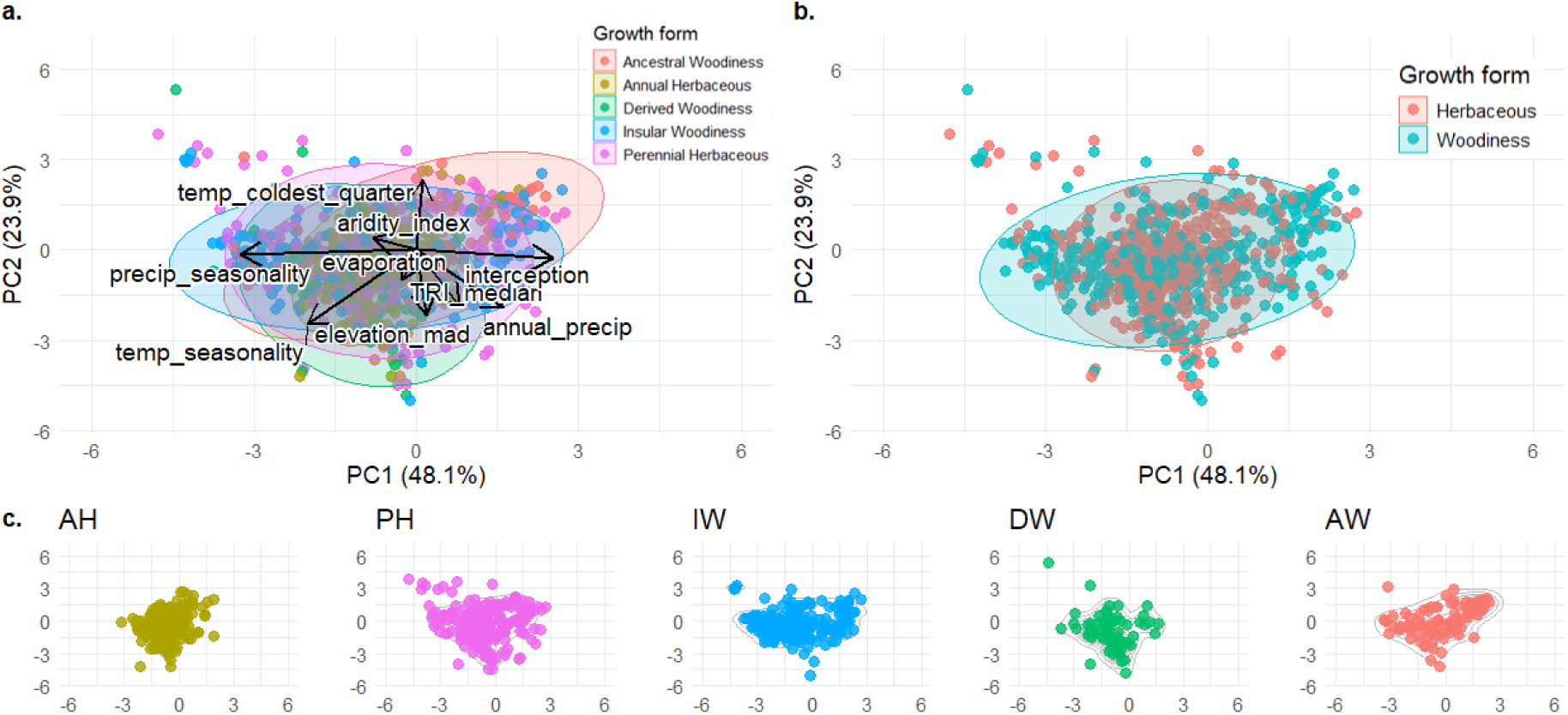
PCA results of the lenient sampling approach. Formatted in the same way as Figure 2.

