## Supplementary Methods M1 for "Climatic seasonality and topographic complexity shape plant growth form distributions in the Canary Islands"

[section numbers correspond with sections in the main text]

**2.1 Locality data extraction**

2.1.2. GBIF extraction & Locality data filtering

Occurrence data was filtered in four steps. First, we filtered locality data using the ’clean_coordinates’ function of the ’CoordinateCleaner’ package (Zizka et al., 2019). After evaluating the results of these initial cleaning criteria, further filtering steps were implemented to remove occurrences that still displayed erroneous patterns. We implemented a custom function to check whether the occurrence coordinates were assigned to islands where the respective plant species are known to natively occur (following Beierkuhnlein et al., 2021). Coordinates on smaller islets and islands were categorised under the name of the island to which they were closest in proximity (e.g. coordinates on La Graciosa and Alegranza were extracted as occurring on Lanzarote).

We also wrote a custom function to remove reprojected occurrences (i.e., from raster to vector data). Due to the reprojection, these occurrences no longer corresponded to exact localities where the species occurs, but rather to the central point in the raster grid prior to reprojection. As such, including these occurrences would bias our dataset due to the inaccuracies stemming from reprojection. These occurrences were removed by filtering out occurrences for which the code ’cdrep’ was assigned in the extracted GBIF data. After this, we performed a final filtering step based on the reported coordinate uncertainty; occurrences with a coordinate uncertainty above a certain threshold were filtered out. To do this, three different sampling regimes were used (see main text).

**Table SM1. Criteria used to filter GBIF-extracted occurrence data**. In total, nine functions from the ’CoordinateCleaner’ and three custom functions are used.

| Function | Description |
| --- | --- |
| CoordinateCleaner | |
| cc_cen | coordinates near island centroids |
| cc_equ | coordinates have equal (absolute) longitude/latitude |
| cc_gbif | coordinates near GBIF headquarters |
| cc_inst | coordinates near botanical institutions |
| cc_zero | both coordinates are 0 |
| cc_coun | country-label does not match coordinates |
| cc_sea | coordinates are located in open waters |
| cc_cap | coordinates near island capitals |
| cc_dupl | occurrences have equal coordinates |
| Custom functions | |
| island_occurrence | coordinates occur on non-native island |
| cdrep_issue | ‘cdrep’ issue is reported for occurrence |
| uncertainty | coordinate uncertainty is higher than pre-defined limit |

**2.2 Environmental dataset compilation**

2.2.1. Variable extraction

We selected nine environmental variables from the CanaryClim v1.0 dataset (Table 2; Patino et al., 2023), comprising both topographic and climatic variables. To investigate the evolution of insular woodiness on the Canary Islands, we included variables associated with its evolution: diurnal and seasonal temperature variability, temperature of the coldest quarter, and precipitation seasonality (calculated as the coefficient of variation of the monthly precipitation estimates) to assess the aseasonal favourable climate hypothesis (Carlquist, 1974); and the air temperature of the driest quarter, annual precipitation amount, and the precipitation in the driest quarter to assess the drought hypothesis (Zizka et al., 2022).

Additionally, data on evapotranspiration and interception (hereafter referred to as evapotranspiration) were included (FAO, 2020). Using the evapotranspiration and precipitation data, an (adjusted Thornthwaite) aridity index was calculated (Herschy et al., 1998), quantifying the ratio of water entering (through precipitation) to water leaving (through evapotranspiration) the environment via plant species, as described in equation (1):

$AI_{s} = \frac{P_{s}}{ET_{s}}$

where, for species 𝑠, 𝐴𝐼𝑠 is the aridity index, 𝑃𝑠 is the median annual precipitation (mm/year), and 𝐸𝑇𝑠 is the median annual actual evapotranspiration and interception (mm/year).

Furthermore, we used the Digital Elevation Model (DEM) of the CanaryClim dataset to extract the elevation for each of the filtered occurrences. Elevation can serve as a proxy for multiple different parameters related to plant growth, such as the amount of solar radiation, air temperature, and precipitation (Körner, 2007). Also, topographic heterogeneity has been recognised as a key variable linked to plant species richness (Liu et al., 2019; Nürk et al., 2020; Chang et al., 2023). As such, we included a Terrain Ruggedness Index (TRI), calculated following the methodology of Riley et al. (1999). TRI quantifies topographic heterogeneity by considering the difference in elevation between a cell in a grid and the neighbouring cells. The TRI raster was calculated using the ’tri’ function in the ’spatialEco’ package (Evans & Murphy, 2023), using a value of 3 as the scale window parameter and the CanaryClim DEM as input.

2.2.3. Growth form determination

We determined the growth form of the species through several methods. First, for IW lineages, we used IW species lists from Lens et al. (2013) and Zizka et al. (2022). For DW and AW lineages, we used Hooft van Huysduynen et al. (2021) and an unpublished dataset by F. Lens. If no information on growth form was found for a species in these studies, the protologue of the species was used, primarily accessed through the International Plant Names Index (IPNI, 2024). In case the protologue could not be found or did not resolve the growth form, a literature search was performed using Google Scholar and the Web of Science (WoS) for the species name and a reference to the growth form (e.g. ’woody’, ’woodiness’, or ’herbaceous’). If no information on growth form was available through the aforementioned steps, information from herbarium sheets, pictures of herbarium sheets, or individuals in the field were used. Lastly, if a species was identified as woody and not in any of the aforementioned studies, the available phylogenetic literature was used to reconstruct the evolutionary history of the woodiness in the clade. The list of all species with the assigned growth forms is provided in Table S1.

**2.3 Environmental data analysis**

2.3.2. Phylogenetic Data, Phylo-PCA, & Phylo-ANOVA

To construct a phylogeny for our analyses, we used V.PhyloMaker2 (Jin and Qian, 2022), a software for generating plant phylogenies based on the vascular plant supertree of Smith and Brown (2018) and the pteridophyte phylogeny of Zanne et al. (2014). For each sampling regime, we produced a separate phylogeny using a list of all the species which remained after locality data filtering, with the ‘GBOTB.extended.TPL’ tree selected, with tree and nodes arguments set to default. All species in our input list were recovered in the phylogeny. This phylogeny can also be replaced by an input tree (newick file) from the user if plants are not the group of interest.

To visualise the major axes of variation across the environmental variables, while accounting for the shared evolutionary history among taxa, we performed a phylogenetic principal component analysis (hereafter ‘phylo-PCA’) as described by Revell (2009; Figure 2), using the ‘phyl.pca’ function from the ‘phytools’ package (Revell, 2024). Before the PCA analysis, the data were centred and scaled to ensure equal contribution to the model. Additionally, we colour-coded the points based on their growth form.

To test for differences in the environmental variables across the five different growth forms, while accounting for their shared evolutionary history, we used phylogenetic Analysis of Variance (phylo-ANOVA; Figure 3). The ‘phylANOVA’ function in the ‘phytools’ R package (Revell, 2024) was used, which simulates trait evolution under a Brownian motion model along the provided phylogenetic tree to generate a null distribution of F-statistics (Garland Jr et al., 1993). The significance of group differences was assessed via 1,000 simulations, and p-values were computed as the proportion of simulated F-values exceeding the observed value. P-values were adjusted for multiple comparisons using the correction procedure by Holm (1979).
